# Extensive length and homology dependent chimerism in pool-packaged AAV libraries

**DOI:** 10.1101/2025.01.14.632594

**Authors:** Jean-Benoît Lalanne, John K. Mich, Chau Huynh, Avery C. Hunker, Troy A. McDiarmid, Boaz P. Levi, Jonathan T. Ting, Jay Shendure

**Affiliations:** Department of Genome Sciences, University of Washington, Seattle, WA, USA; Département de Biochimie et Médecine Moléculaire, Université de Montréal, Montréal, QC, Canada; Allen Institute for Brain Science, Seattle, WA, USA; Seattle Hub for Synthetic Biology, Seattle, WA, USA; Brotman Baty Institute for Precision Medicine, Seattle, WA, USA; Howard Hughes Medical Institute, Seattle, WA, USA; Allen Discovery Center for Cell Lineage Tracing, Seattle, WA, USA

## Abstract

Adeno-associated viruses (AAVs) have emerged as the foremost gene therapy delivery vehicles due to their versatility, durability, and safety profile. Here we demonstrate extensive chimerism, manifesting as pervasive barcode swapping, among complex AAV libraries that are packaged as a pool. The observed chimerism is length- and homology-dependent but capsid-independent, in some cases affecting the majority of packaged AAV genomes. These results have implications for the design and deployment of functional AAV libraries in both research and clinical settings.

## Main Text

Multiplexed functional genomic screens often employ linked components within a cargo, for example single-cell CRISPR screens^1,2^ (sgRNA and barcode) or massively parallel reporter assays^3^ (MPRAs, regulatory element and barcode). In these, any level of decoupling of expected pairings ultimately degrades signal quality. A prominent example is the frequent recombination seen in lentiviral cargo packaging. Such recombination, which is length and homology dependent and results from lowly processive reverse transcription and template switching during replication^4^, stymied early single-cell functional genomics efforts using this delivery strategy^5^.

In contrast, pervasive chimeric rearrangements of AAV genetic material have not been described to our knowledge. Yet, recent studies of long-read sequenced AAV-packaged DNA have revealed unexpected DNA arrangements^6,7^. In parallel, high levels of noise and limited dynamic range are commonly observed in barcoded MPRA experiments using AAVs^8–12^ (see also Hunker et al., co-submitted). These hint at possible unknown complexities during AAV packaging.

To explicitly test for chimera formation during AAV packaging, we performed a barcode swap experiment. Specifically, we constructed a series of complex AAV libraries with uniquely associated pairs of barcodes separated by different inserts and flanked by AAV2 inverted terminal repeats (ITRs) (**Fig. 1a-b**). The six inserts consisted of three lengths (short: ∼130 bp, mid-sized: ∼740 bp, long: ∼2 kb) each with two classes (homologous: identical sequences; non-homologous: size-matched to homologous counterparts and bearing tagmented and narrowly size-selected *E. coli* genomic DNA; **Fig. 1a, S1, S2**). Each insert further had a short internal sequence index for downstream demultiplexing (**Fig. S1d**). The resulting libraries were complex (>5M barcode pairs in parental p146, bottlenecked to 15-45k pairs in p149-p154). Cargoes were size-adjusted with filler sequences to fix the total ITR-to-ITR length to ≈2.3 kb for inserts of different sizes (**Fig. S1c**, but see library p153^×^ below). These libraries were then packaged separately into AAVs (all with PHP.eB, some with AAV2 serotypes, 9 packaging conditions total, **Table S4, Methods**).

**Figure 1:**
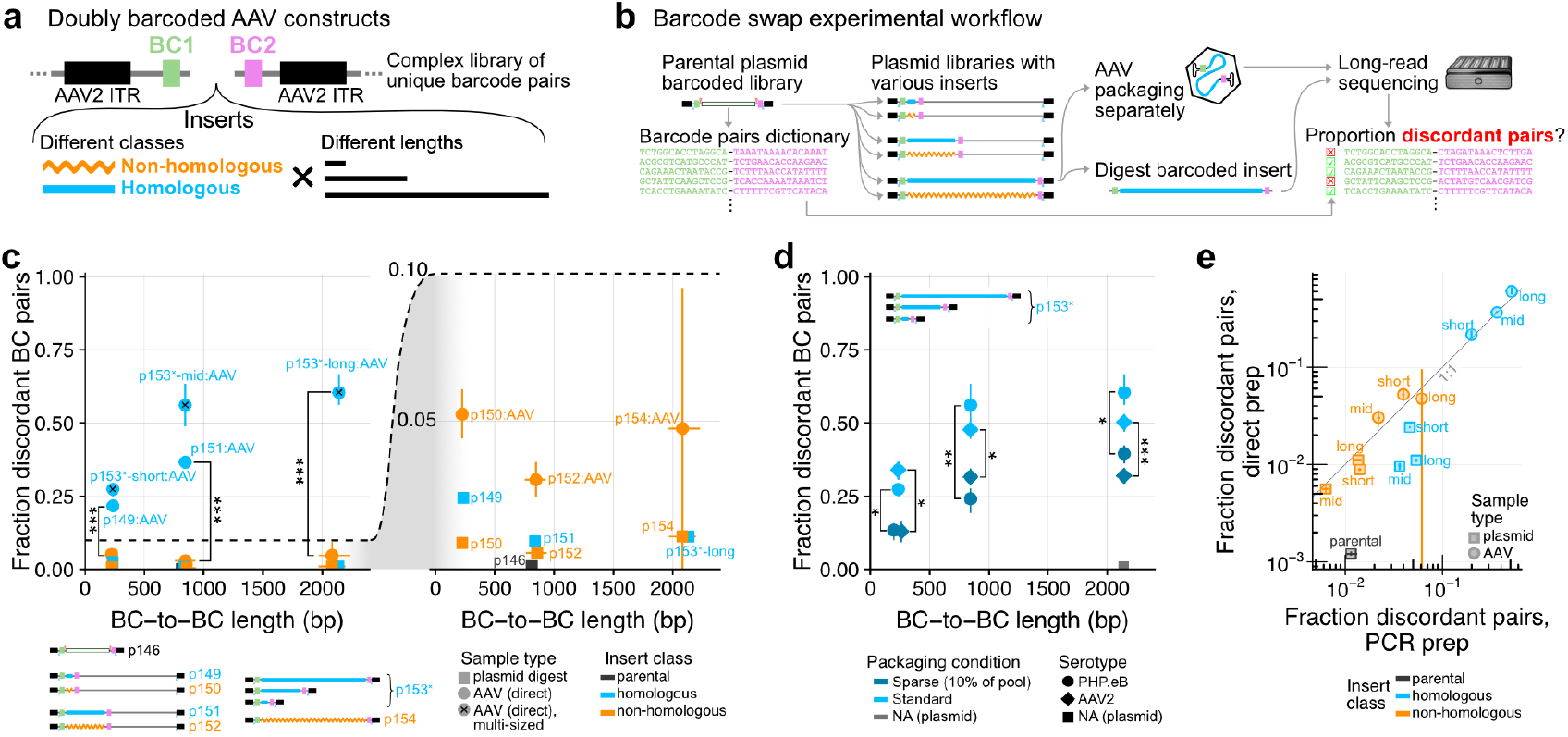
Chimera formation during AAV packaging revealed by barcode swapping experiments. **(a)** A complex doubly barcoded cloning dock with an *a priori* determined valid barcode pairs dictionary serves as the starting point to clone libraries of barcoded inserts varying length and class (homologous & non-homologous) within an AAV cargo. **(b)** Barcoded libraries were separately packaged in AAVs, and submitted for direct long-read sequencing, in parallel with corresponding insert size-selected from the plasmid digest. **(c)** Quantification of the fraction of discordant barcode pairs as a function of the full-length BC-to-BC average size. Each point corresponds to swap quantification from 6 separate libraries ([short, mid, long] × [homologous, non-homologous]) for both plasmid-derived (square) and AAV-derived (circle) material (n=1 replicate per library). For each datapoint, we analyzed full length BC-to-BC reads passing quality control filters and having separately valid BC1 and BC2 (**Fig. S3a, S4a**). Right panel corresponds to a zoomed inset showing the y axis range from 0 to 0.1. Quantifications derived from library p153^×^ corresponding to the pool of multiply-sized inserts in a single AAV-packaged sample are marked by an ×. Vertical error-bars correspond to 20^th^ to 80^th^ percentiles from bootstrap resampling. Horizontal error-bars to the 10th to 90th percentile of the BC-to-BC length distribution from plasmid digest inserts. Swaps are significantly higher (one-sided bootstrap FDR: ***<10^−5^) for the homologous vs. their respective size-matched non-homologous libraries. **(d)** Same as (c), but with additional packaging conditions with library p153^×^ corresponding to AAV2 serotype and sparse co-packaging with carrier DNA. With both PHP.eB and AAV2, and for the three sizes of inserts, sparse packaging reduces swapping significantly (one-sided bootstrap FDR: *<0.005, **<0.0005, ***<10^−5^) albeit incompletely. **(e)** Comparison of the fraction of discordant barcode pairs from PCR-derived libraries (x-axis) vs. direct [same data as panel (c)] (y-axis), see also **Fig. S5b**. AAV samples only include the PHP.eB serotype with standard packaging conditions. Errorbars correspond to 20^th^ to 80^th^ percentiles from bootstrap resampling of reads as in panel (c).

To assess for chimeras, defined as discordant barcode pairs within a single read, we performed long-read sequencing of the barcoded inserts using PCR-free library preparation, from both sized-selected digested plasmids (“zero-swap” controls) and AAV-packaged DNA, on the Oxford Nanopore platform (Plasmidsaurus). Focusing on full length BC-to-BC reads bearing all signposts in their proper positions and exact but separate BC1 and BC2 matches (**Fig. S1d, S3a-b**), we measured what fraction of reads showed concordant versus discordant BC1-BC2 pairs in each sample (**Fig. 1c, S3e, S4d**). As expected, the parental p146 BC1-BC2 plasmid library backbone exhibited near-complete concordance (0.1% discordant pairs, **Fig. 1c**). Also, the zero-swap inserts digested from plasmid library controls p149-p154 showed low but slightly increased discordance (<2.5%, right inset **Fig. 1c**), suggesting non-zero chimerism generated in the cloning process (the highest discordance library being in the short-homologous p149, hinting at possible recombination during Gibson assembly).

Remarkably however, homologous insert AAV-packaged libraries exhibited dramatically increased rates of discordance, ranging from ≈20% for the short inserts to >60% for the long inserts (**Fig. 1c**, left). In stark contrast, non-homologous libraries displayed largely concordant barcode pairs independent of length (≤6% discordance, significantly lower than size-matched homologous libraries, FDR<10^−5^ by bootstrap analysis). We note that due to the library construction process (tagmentation followed by PCR), even the non-homologous inserts shared short regions of homology corresponding to the Tn5 adapters at both ends (33+34=67 bp), which could contribute to the low but non-zero rates of chimerism in these. Further, a similar level of chimerism was observed in an orthogonally prepared low-complexity library of pool-packaged barcoded enhancer AAV constructs (Hunker et al., co-submitted). Taken together, these results indicate extensive molecular chimerism forming during AAV packaging, in some cases representing the majority of species, and suggest that the rate of chimera formation is dependent on the length of intervening homologous sequence.

To assess whether the observed chimerism was exclusive to serotype PHP.eB, library p153^×^ was also packaged with AAV2, revealing a similar level of barcode swaps (**Fig. 1d**). We also attempted to minimize chimerism by transfecting at sparse conditions (10% barcoded library p153^×^ with 90% carrier DNA). Sparse packaging did decrease chimerism as expected (bootstrap FDR<0.005 in all instances), but could not completely eliminate this phenomenon (**Fig. 1d**). These results suggest that AAV chimeras form in a variety of capsids and packaging conditions.

Three technical points deserve note. First, one of our libraries, p153, while intended to exclusively harbour long homologous inserts, also contained a sizeable proportion of short and mid-sized homologous inserts in the AAV-packaged ONT data due to cloning history (short: 8 to 49%, mid-sized: 10 to 23%, across the four samples), which could be identified due to their internal insert index and shorter total read lengths (**Fig. S4e**). Quantifications from the resulting multi-sized library, denoted as p153^×^, are marked with × in **Fig. 1c** (all points in **Fig. 1d**). Shorter insert elements in p153^×^ were present at low proportion in the starting plasmid library (not detected in whole plasmid sequencing verification & barely visible on gel but clearly visible upon PCR amplification, **Fig. S4g-i**), but were likely selectively enriched due to their shorter sizes during AAV packaging. All other libraries were overwhelmingly comprised of the expected inserts (>99% apart from generally low level of empty parental carryover; mid-sized and long non-homologous displayed higher proportion of parental p146 sequences, at 19% and 66% respectively, possibly enriched by the annealing step of the ONT library preparation, see below).

Second, AAV-packaged long inserts (p153^×^-long and p154) showed a high proportion of reads with shorter than expected BC-to-BC inserts (**Fig. S4d**, 14 to 67% in p153^×^, 85% in p154). These off-products (not included in the quantification shown in **Fig. 1c-d**) were associated with even lower rates of barcode pair concordance, irrespective of the homologous or non-homologous nature of their inserts (**Fig. S4e**). Inserts from these shorter off-products displayed a high proportion of complex composite multi-segment alignments (**Table S2**, Fisher’s exact p<10^−4^ in all instances), in-line with previous evidence of complex structural variants in AAV-packaged DNA^6,7^.

Third, some of the reads, despite having the full BC-to-BC length, did not span the full ITR-to-ITR length. This phenomenon was seen not only in the AAV-packaged samples but also in the size-selected digested plasmid inserts (c.f., **Fig. S3f** vs. **S4e**) which were overwhelmingly ≈2.2 kb upon submission for long-read sequencing. Therefore, we tentatively attribute these to downstream technical aspects of the ONT sequencing and not to biological processes during AAV packaging. Consistently, these incomplete ITR-to-ITR reads (but still with full BC-to-BC lengths) had similar rates of barcode swapping (Fisher’s exact p>0.35 in all instances, **Table S3**).

What alternative explanations would account for the observed chimerism? The AAV sequencing library preparation from Plasmidsaurus, based on a recent protocol^13^, relies on annealing of the ssAAV DNA followed by end-repair and adapter ligation. End-repair can in principle induce a single polymerization event, such that incomplete products with an insert-internal 3’ end could prime homologous counterparts, leading to technically induced chimeras. This is, however, unlikely, as spurious priming-competent products with internal 3’ ends in principle should not arise since AAVs are packaged from their 3’ ITR^14,15^. Further, truncated/incomplete AAV genomes^16^ would need to make up a substantial proportion of the packaged material commensurate with the observed barcode swapping frequency, but are a rare class of intermediates compared to snapback products^17^ (partial genomes flanked by ITRs on both ends). Finally, AAV-dimers generated from ITR priming, which serve as an internal control for this effect, are rarely observed in ONT libraries generated using this method (N. Kamps-Hughes, personal communication).

To confirm that barcode-swapping chimeras were not a technical consequence of the direct ONT sample preparation from AAVs, we also generated double-stranded DNA libraries of barcoded inserts by PCR templated from both plasmids and purified AAV-packaged genomes. Of note, PCR libraries also avoided a possible confounder of libraries composed of large segments of non-homology (p152 and p154), which might have hindered the annealing step in the direct AAV ONT preparation. Plasmid templates were diluted to have a similar number of PCR cycles total compared to the AAV genomes templates (n=16-20 cycles for AAV samples, n=18 cycles from plasmid). Long-read sequencing (Plasmidsaurus) and processing of the PCR-generated libraries as before revealed near-quantitative agreement in the level of swaps compared to the PCR-free samples (**Fig. 1e, S5**). The main difference observed was a modest increase in swaps for plasmid samples originating from homologous-insert (blue squares in **Fig. 1e**, from 1-2% [direct] to 3-5% [PCR] discordant pairs), in line with the low-level of chimerism known to originate from PCR^18^. This PCR chimerism was nevertheless about one order of magnitude lower than that observed in AAV-derived samples. Further, this result is consistent with nearly nonexistent chimerism (<2.5% swaps) in a sample composed of separately packaged plasmids pooled prior to ONT processing (Hunker et al, co-submitted). In sum, these data strongly support the interpretation that the observed chimerism was not introduced by the AAV ONT library preparation process.

A critical aspect of *in vivo* functional genomics is high-fidelity delivery of functional payloads to cells, and the scale of these experiments has been growing^19^. AAV vectors have known limitations in terms of the size of their DNA payload, but to our knowledge have not previously been reported to undergo high frequency rearrangement of their genetic content upon packaging. This is unlike the well-established case of lentiviral vectors, which have long been known to recombine due to template switching during packaging, thereby dramatically unlinking their genetic components^4,5,20–23^ in a length and homology dependent manner. Our results document widespread chimerism within AAV particles as well. This chimerism is likely a substantial contributor to the high levels of noise frequently observed in multiplexed barcoded AAV packagings by ourselves and others^8–12,24^ (see also Hunker et al., co-submitted, which provides evidence of consequences for the interpretation of *in vivo* experiments). The exact mechanism of this chimera formation is presently unknown and beyond the scope of this brief communication, but the dependencies on length, homology, and cotransfection density suggest that chimera formation could form via template switching of partial genomes during rolling hairpin amplification^25^, analogous to that commonly observed in PCR with poorly processive polymerases^18,26,27^. Regardless, this effect can potentially be mitigated^5,22,23^ by limiting the distance between complex elements (e.g., 5’ instead of 3’ barcodes in MPRA assays), decreasing the extent of entirely homologous regions, and/or lowering cotransfection rates in packaging cells when possible (at the expense of titer yield), or dispensing of the need for a barcode altogether where possible (e.g., direct capture of sgRNAs^28^). Future technical improvements in packaging cofactors could also find general solutions to this problem. Taken together these results illuminate a problem in AAV production, which will inform experimental design decisions to improve data quality in projects involving this important gene delivery vehicle.

## Acknowledgements

We thank Nick Donadio, Rana Kutsal, Shannon Khem, and Shenqin Yao for AAV packaging support at the Allen Institute, Nicolas Kamps-Hughes (Plasmidsaurus) for technical comments, and the entire Shendure lab for discussion. We thank Molly Gasperini for critical reading of the manuscript. This work was supported by the National Institutes of Health (R01HG010632 to J.S.). J.-B.L. was supported by the Damon Runyon Foundation (DRG-2435-21) and by a Next Generation Scientist award from the Cancer Research Society of Canada (grant no. 1155581). T.A.M was supported by a Banting Postdoctoral Fellowship from the Natural Sciences and Engineering Research Council of Canada. J.S. is an Investigator of the Howard Hughes Medical Institute. Allen Institute authors wish to thank Paul G. Allen Family Foundation and NIH BRAIN Initiative Armamentarium Grant UF1MH128339 (to J.T.T., B.P.L.) for their support.

## Author contributions

J.-B.L., J.K.M, and A.C.H. conceptualized the barcode swap experiment with input from T.A.M. J.-B.L. and C.H. cloned libraries. J.-B.L. analyzed data. J.-B.L., J.K.M, and J.S. wrote the manuscript. B.P.L., J.T.T., and J.S. supervised the study.

## Competing Interests

J.S. is a scientific advisory board member, consultant and/or co-founder of Cajal Neuroscience, Guardant Health, Maze Therapeutics, Camp4 Therapeutics, Phase Genomics, Adaptive Biotechnologies, Scale Biosciences, Sixth Street Capital, Prime Medicine, Somite Therapeutics and Pacific Biosciences. J.K.M. and B.P.L. are founders of EpiCure Therapeutics, Inc. Other authors declare no competing interests.

## Data availability

All scripts used to analyze data and plasmid maps/amplicon (for p146 BC pair dictionary generation) files have been deposited to Zenodo (10.5281/zenodo.14515777, URL for access). Raw short read (p146 barcode dictionary generation) and long-read (ONT, barcode swap assessment) sequencing data have been submitted to GEO (accession GSE284548, access token: iriheiswvrullcn), together with the following processed files:

Barcode pair dictionary generation:

> Piled-up raw short-reads to pairs (with >1 reads) of barcodes:
>
> *p146_2BC_backbone_S1_BC_pairs_condensed_no1_20241101*.*txt*.*gz*
>
> List of well-represented BC1 and BC2:
>
> *final_set_leftBC1_categories_p146_20241105*.*txt*.*gz*
>
> *final_set_rightBC2_categories_p146_20241105*.*txt*.*gz*
>
> Final valid barcode pairs dictionary table (from parental p146):
>
> *final_paired_BC1_BC2_p146_20241216*.*txt*.*gz*

Processed long-read data:

Table of ONT reads with coherently positioned signposts:

Direct, plasmid (n=887145 reads): *ONT_reads_all_signposts_plasmid_digest_20241216*.*txt*.*gz*

Direct, AAV (n=11294 reads): *ONT_reads_all_signposts_AAV_direct_20241216*.*txt*.*gz*

PCR-derived, plasmids (n=215315 reads): *ONT_reads_all_signposts_plasmid_PCR_20250103*.*txt*.*gz*

PCR-derived, AAV (n=270519 reads): *ONT_reads_all_signposts_AAV_PCR_20250103*.*txt*.*gz*

Table of ONT reads with coherently positioned signposts, demultiplexable, with separately valid BC1/BC2:

Direct, plasmid (n=392391 reads): *ONT_final_valid_BCs_reads_plasmid_digest_20241216*.*txt*.*gz*

Direct, AAV (n=4811 reads): *ONT_final_valid_BCs_reads_AAV_direct_20241216*.*txt*.*gz*

PCR-derived, plasmids (n=113955 reads): *ONT_final_valid_BCs_reads_AAV_PCR_20250103*.*txt*.*gz*

PCR-derived, AAV (n=123969 reads): *ONT_final_valid_BCs_reads_AAV_PCR_20250103*.*txt*.*gz*

## Supplementary files

Detailed methods p. 16 to 22 in this document. Associated data tables (in addition to tables found in **Fig. S2-S5**):

**Table S1**: List of plasmids and oligos used.

**Table S2:** Insert alignment metadata and read counts stratified by full/non-full BC-to-BC length.

**Table S3:** Table of read counts of barcode swap stratified by full/partial ITR-to-ITR length.

**Table S4:** AAV packaging metadata.

---

**Figure S1:**
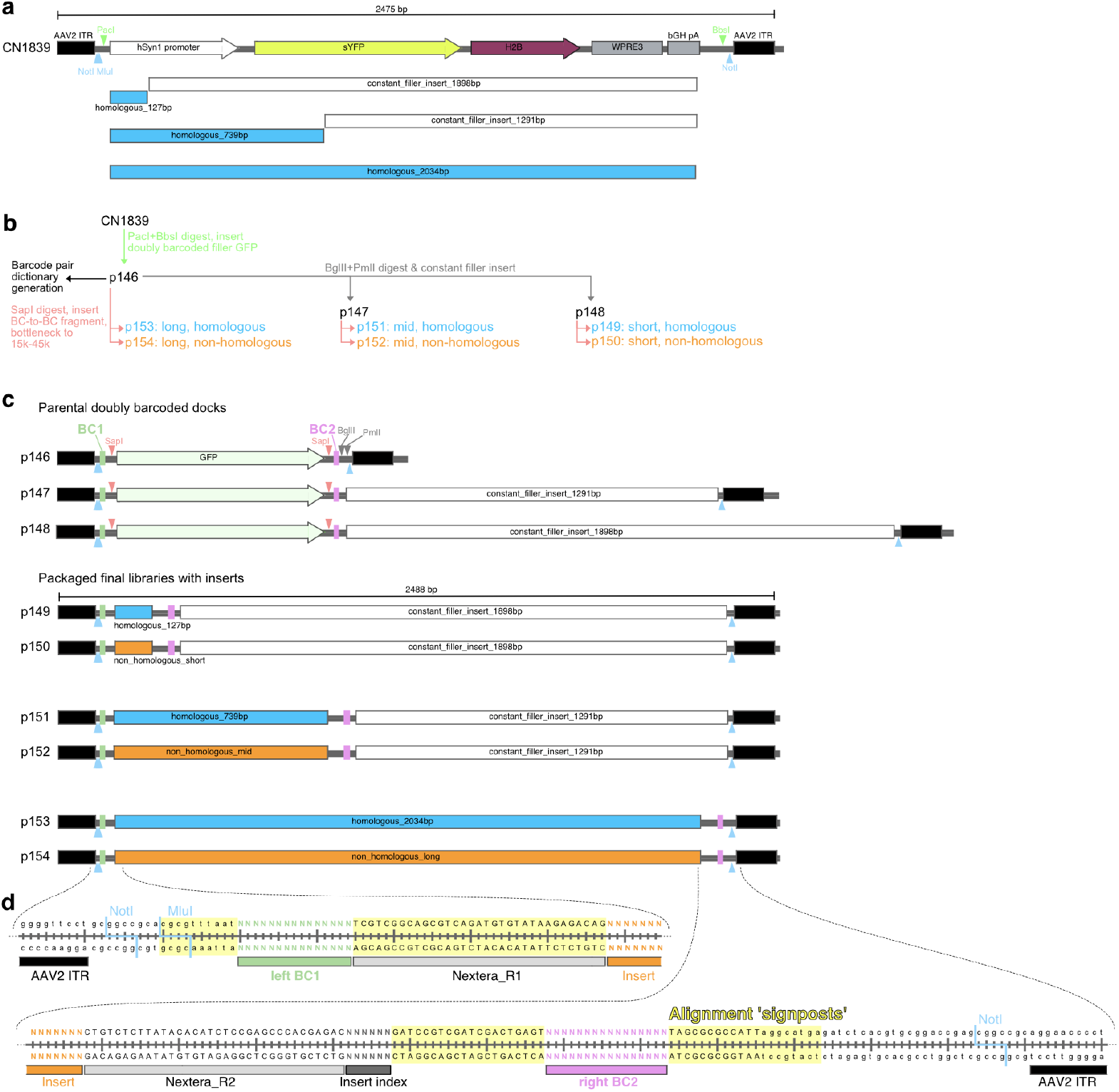
Barcode pairs AAV constructs and cloning strategy. **(a)** Schematic of components of plasmid CN1839 (Addgene#163509) between the AAV2 ITRs. Segments under the map highlight constant regions (cloned by PCR) used both as homologous library inserts (blue) and filler sequences to fix the ITR-to-ITR length (white). These segments are shown panel (c) below. Positions of PacI and BbsI restriction sites are marked by green carets, NotI and MluI sites by pale blue carets. **(b)** Molecular cloning scheme used. First complex BC1-BC2 parental dock with GFP stuffer p146 was cloned, and served as template for cloning secondary docks p147 and p148 with filler sequences. Complex library p146 was used as template for barcode dictionary generation. All parental docks were digested with SapI to liberate the GFP, which was replaced by respective internally indexed inserts to generate the final series p149-p154 (which were all bottlenecked to a target of 20k transformants). **(c)** At scale schematics of ITR-to-ITR components of cloned parental and insert-containing libraries. Position of SapI restriction sites are marked by pale red carets, BglII and PmlI sites by grey carets. **(d)** Sequences surrounding the two barcodes highlighting alignment signposts (yellow) used to create local position reference frames around barcodes in the long-read data. The ‘Insert index’ is shown as Ns, but is fixed and different for each type of insert (not degenerate). Constant Nextera handles between the barcodes (which constitute short homologous regions even in the non-homologous libraries) are shown.

**Figure S2:**
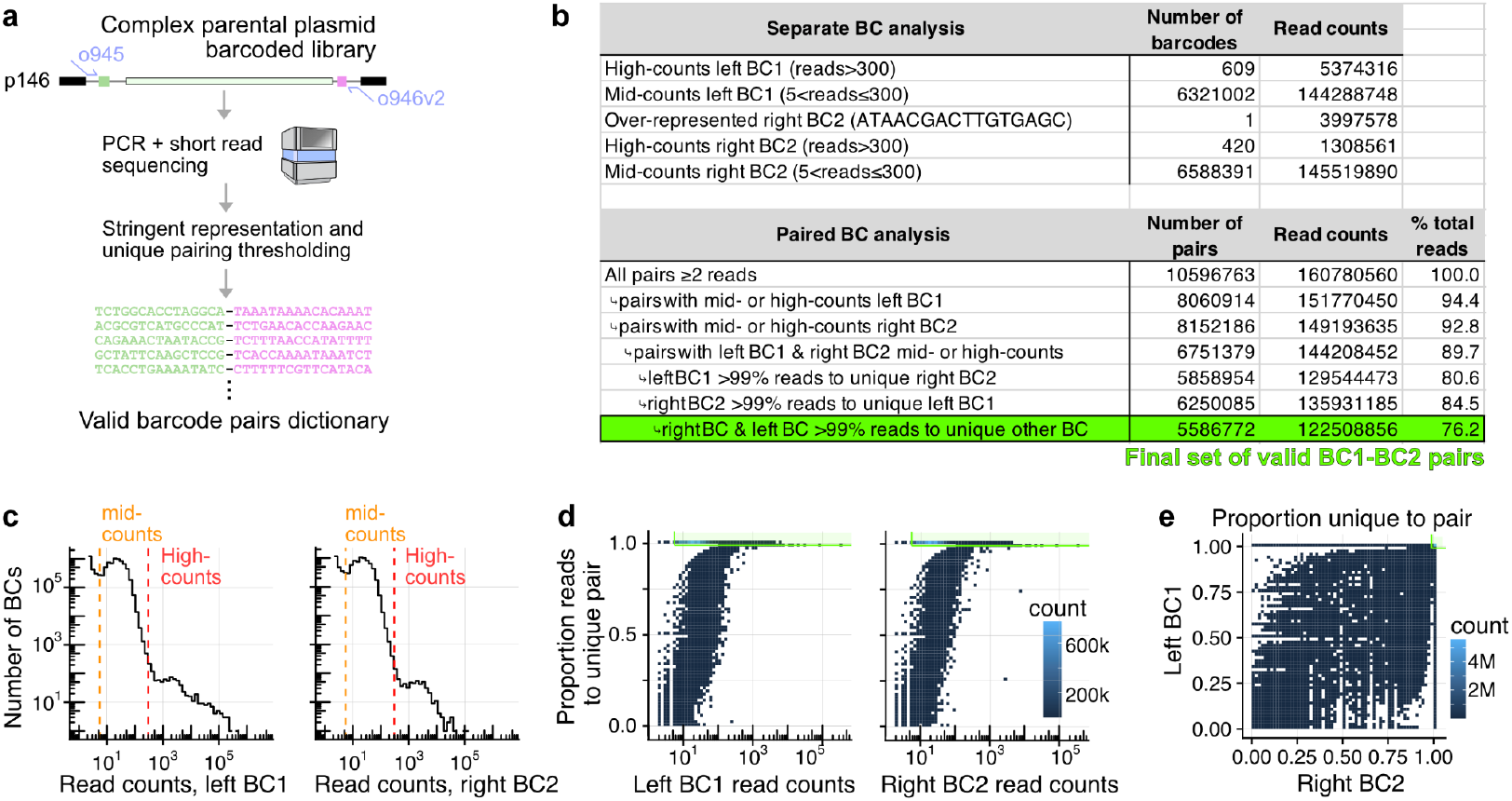
Barcode pairs dictionary generation with paired-end short read sequencing. **(a)** Schematic of procedure to generate the BC1-BC2 pairs dictionary. Parental plasmid dock p146 was used as template for PCR (primers o945+o946v2) to append Illumina P5 and P7 handles, and sequenced on NS2000 with paired-end sequencing to retrieve barcode pair representation. **(b)** Top: Table of number of barcodes and reads for the different categories shown in panel (c). Bottom: Table of number of barcode pairs, reads, and proportion displaying retention at every filtering step (i.e., both barcodes present in the well-represented set and uniquely paired [>99% reads] with a single other barcode). **(c)** Read count distribution by summing only on respective barcodes (not pairs) for BC1 (left) and BC2 (right). Dashed lines indicate cut-offs used for the mid- and high-counts classes of BCs. **(d)** Two-dimensional distributions showing the proportion of reads to the BC arising from the pair on y-axis (BC1 left panel, BC2 right panel) vs. the total read count to the given pair (x-axis). Retained pairs are within the green boundary (>5 reads and >99% unique proportion). **(e)** Similar to panel (d), but now showing unique pairing proportions for BC1 (y-axis) vs. BC2 (x-axis). Retained pairs are in the top right corner within the green boundaries (>99% uniqueness to both).

**Figure S3:**
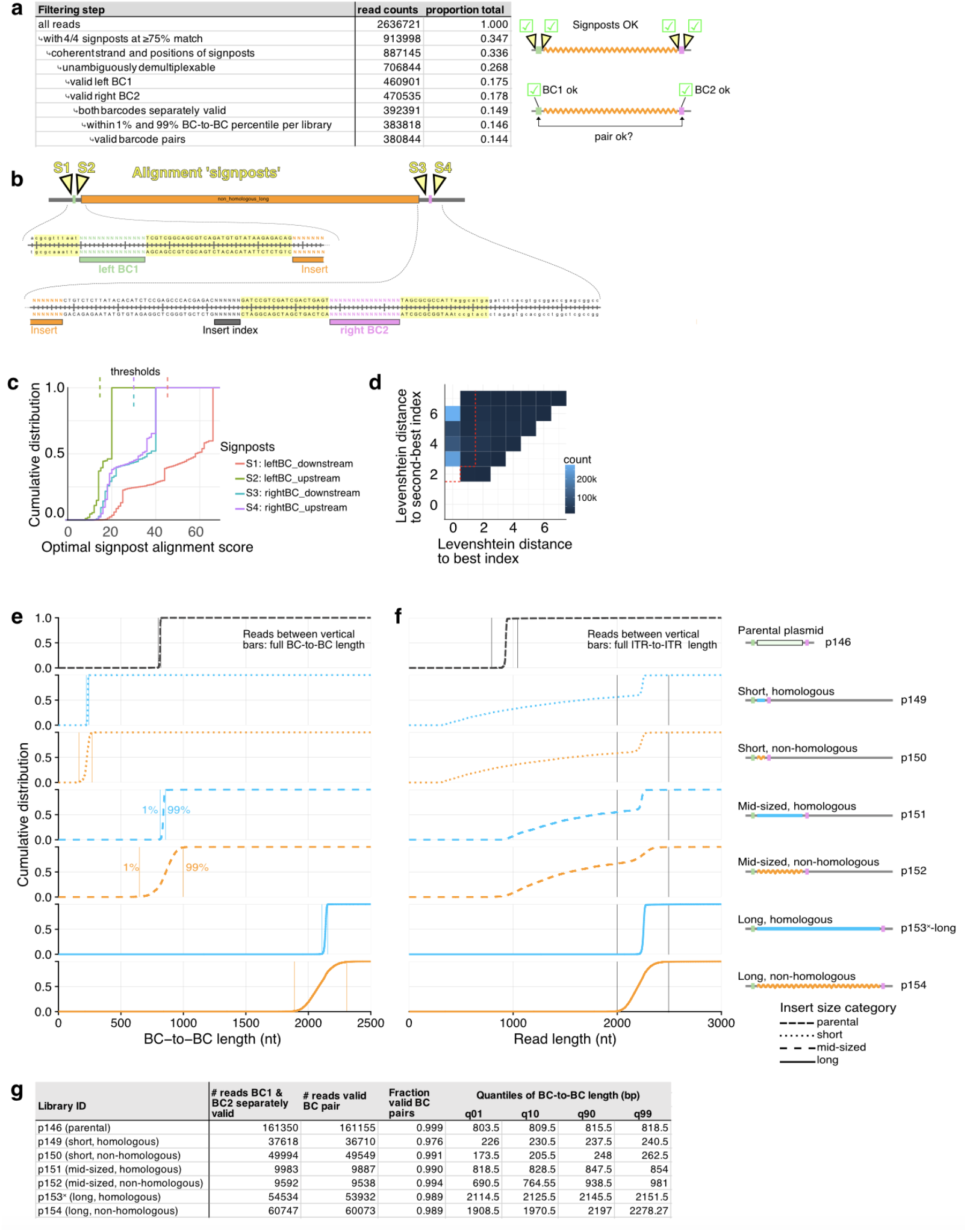
Quality control and statistics of long-read data: size-selected plasmid digests. **(a)** Table showing long-read counts retention at different filtering steps in our pipeline (schematic at right indicate filter nature, e.g., presence of 4/4 signpost sequences and (separate) exact separate matches to BC1 and BC2 from our dictionary. **(b)** Repeat of **Fig. S1d** re-indicating signpost sequences used to parse the long-read for its insert, BC1, BC2, and insert index. **(c)** Cumulative distribution of alignment scores to the signposts. Thresholds for detection (75% match) are shown as dashed lines at top. **(d)** Two-dimensional histogram showing the Levenshtein distance to the best and second best matches to insert indices. Decision boundary to deem insert demultiplexing as unambiguous is shown by the red dashed line. **(e)** Cumulative distribution of the BC-to-BC length (from middle of both barcodes) as determined by the detected signpost positions from the reads passing signpost quality control steps and with separate exact matches to BC1 and BC2. Line type is related to insert size category (parental, short, mid-sized, long), color to insert class (black: parental, light blue: homologous, orange: non-homologous). Vertical lines indicate 1% and 99% percentiles of the length distributions respectively (used to consider a long-read in the AAV-packaged data as ‘full BC-to-BC length’). **(f)** Similar to panel (e), but for total read length. Vertical lines used to consider whether the read is full length or not (parental: 800 to 1050 nt, all other: 2000 to 2500 nt) for analysis in **Table S3. (g)** Table of counts of valid reads stratified by demultiplexed libraries (based on insert index), tallying proportion of reads with valid barcode pairs. Quantiles (1%, 10%, 90%, 99%) of the BC-to-BC length across the libraries is also shown.

**Figure S4:**
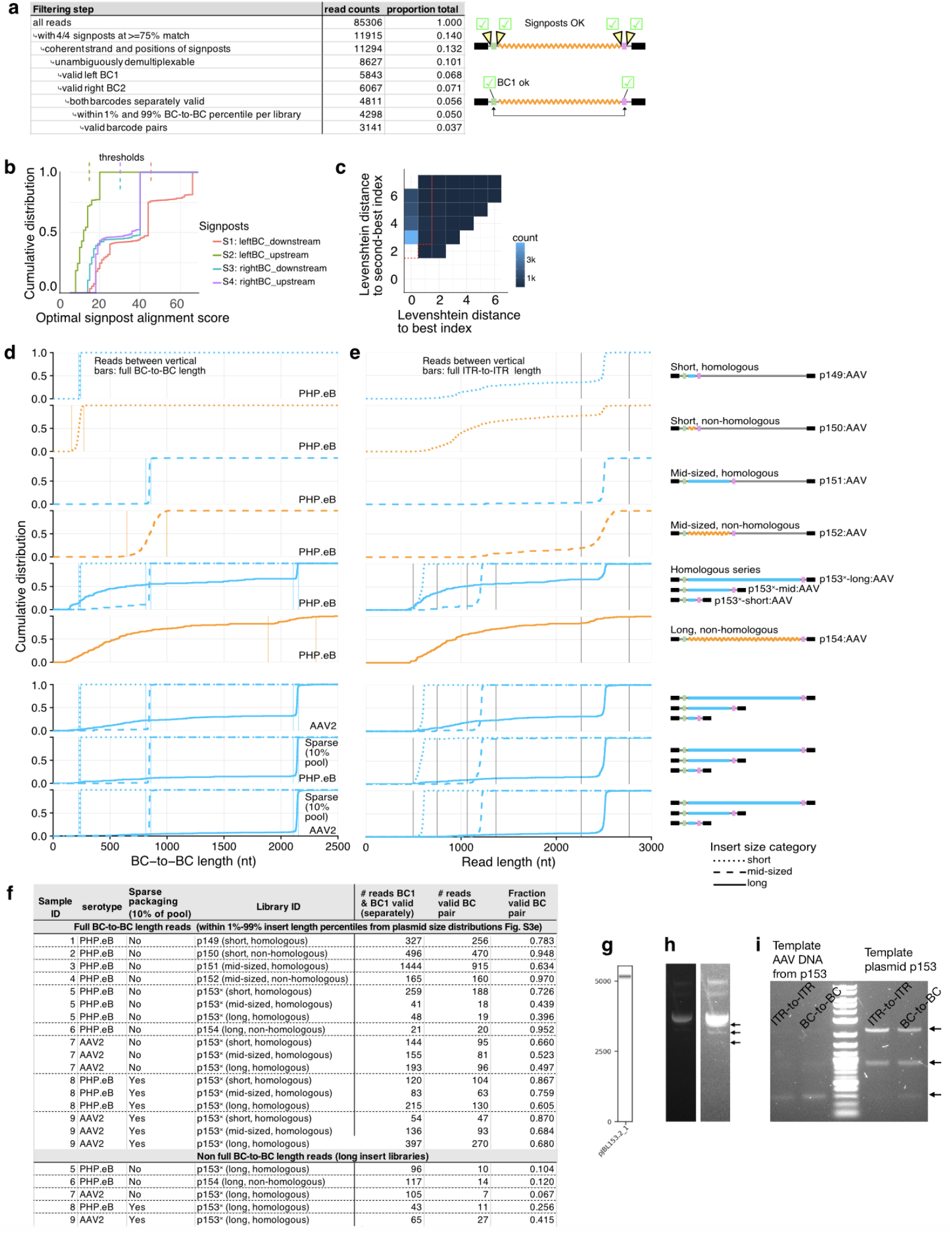
Quality control and statistics of long-read data: direct AAV. Similar to **Fig. S3** with related modifications. **(a)** Table showing long-read counts retention at different filtering steps in our pipeline (schematic at right indicate filter nature), e.g., presence of 4/4 signpost sequences and (separate) exact matches to BC1 and BC2 from our dictionary. **(b)** Cumulative distribution of alignment scores to the signposts. Thresholds for detection (75% match) are shown as dashed lines at top. **(c)** Two-dimensional histogram showing the Levenshtein distance to the best and second best matches to insert indices. Decision boundary to deem insert demultiplexing as unambiguous is shown by the red dashed line. **(d)** Cumulative distribution of the BC-to-BC length (from middle of both barcodes) as determined by the detected signpost positions from the reads passing signpost quality control steps and with separate exact matches to BC1 and BC2. Each panel comes from a separate AAV packaging sample, and was indexed separately (Plasmidsaurus) for ONT sequencing. For p153^×^ samples, different inserts were demultiplexed using the internal insert index. Line type is related to insert size category (parental, short, mid-sized, long), color to insert class (black: parental, light blue: homologous, orange: non-homologous). Vertical lines indicate 1% and 99% percentiles from the plasmid libraries (same as **Fig. S3e**) used to call a read ‘full BC-to-BC length’. Note the substantial fraction of reads with shorter than expected lengths for the long insert libraries. **(e)** Similar to panel (d), but for total read length. Vertical lines used to consider whether the read is full ITR-to-ITR or not (p153^×^-short: 500 to 750 nt, p153^×^-mid: 1100 to 1400 nt, all others: 2250 to 2750 nt) for analysis of **Table S3. (f)** Table of counts of valid reads stratified by demultiplexed libraries (based on insert index), tallying proportion of reads with valid barcode pairs. Quantiles (1%, 10%, 90%, 99%) of the BC-to-BC length across the libraries is also shown. **(g)** Plasmidsaurus (order ID L9RV6B) virtual gel for confirmation of p153, detecting only a single product. **(h)** Agarose gel (low and high contrast) of undigested p153, showing evidence of possible lower molecular weight products (arrows). **(i)** PCR from both AAV template from p153 [serotype PHP.eB, standard condition, sample 5] (left) and plasmid (right) with respectively primers o949+o950 (ITR-to-ITR) and o949+o936 (BC-to-BC). We note that band intensities are probably not representative of species abundance due to possible length-bias of amplification. The three products are of the expected sizes for libraries with short, mid-sized, and long inserts respectively, consistent with their detection in the long-read data.

**Figure S5:**
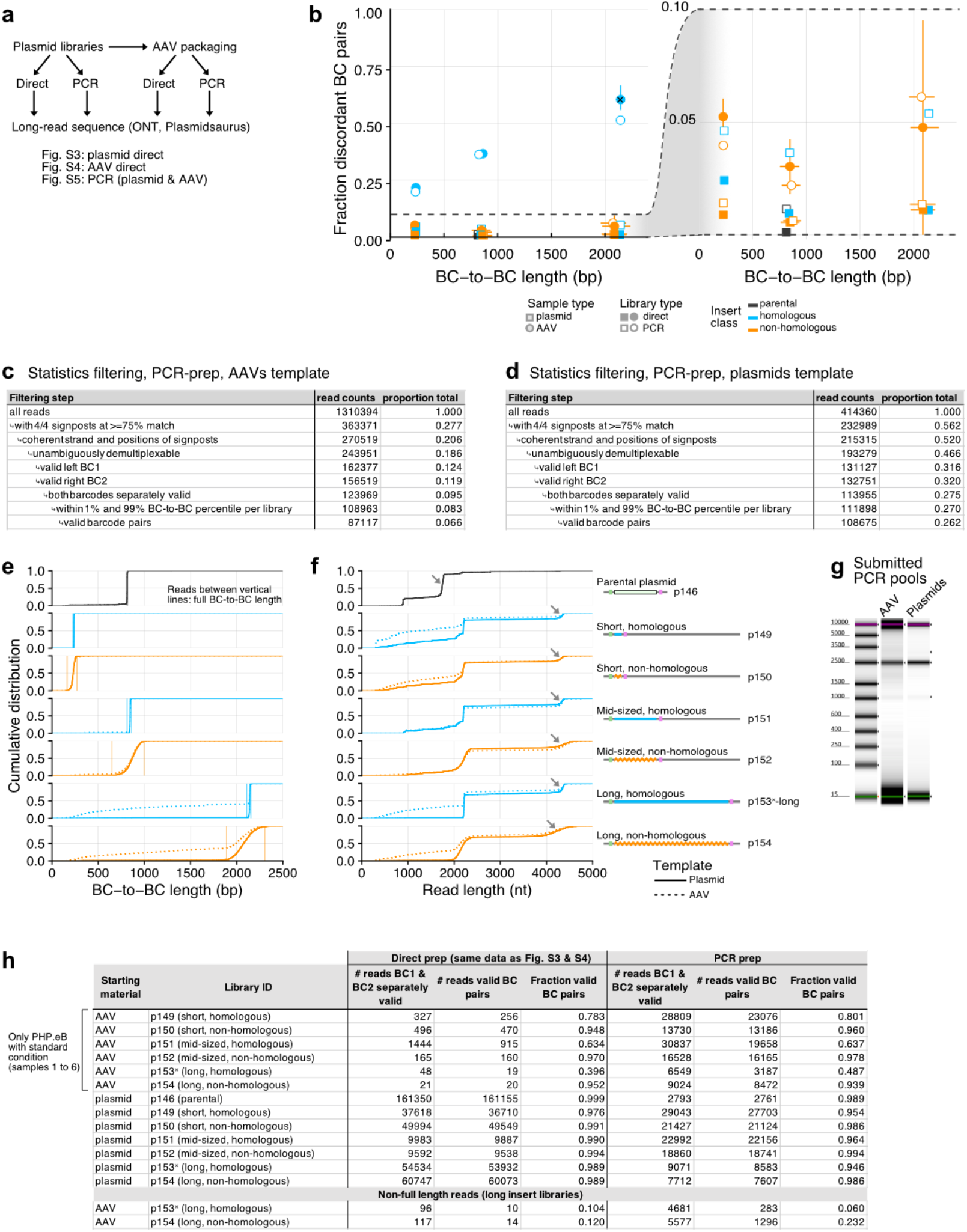
Quality control and statistics of long-read data from PCR libraries. **(a)** Flowchart illustrating the different types of libraries considered (starting material: plasmids & AAV packaged DNA), method (PCR-based vs. direct, i.e., PCR-free, corresponding to digestion & size selection for plasmids, and annealing fpr AAVs, prior to end-repair & adapter ligation). PCR libraries were pooled prior to submission and library identity assigned by demultiplexing based on the internal index (see Fig. S1d). **(b)** Similar to Fig. 1c, now also showing the PCR-derived libraries (open symbols). This data is represented as a comparison between direct/PCR libraries in Fig. 1e. **(c)** and **(d)** Table showing long-read counts retention at different filtering steps in our pipeline for PCR libraries prepared from AAV packaged DNA and plasmids respectively. **(e)** Cumulative distribution of the BC-to-BC length (from middle of both barcodes) as determined by the detected signpost positions from the reads passing signpost quality control steps and with separate exact matches to BC1 and BC2. Line type corresponds to template type (full: plasmid, dashed: AAV), color to insert class (black: parental, light blue: homologous, orange: non-homologous). Vertical lines indicate 1% and 99% percentiles of the length distributions respectively from the plasmid digest (and used to consider a long-read in the AAV-packaged data as ‘full BC-to-BC length’). Panels correspond to different insert libraries. Parental inserts from the AAV sample is not shown as it cannot be reliably attributed to a sample (came from residual empty plasmids packaged from all libraries). **(f)** Similar to panel (e), but for the full read length. Grey arrows indicate ‘dimer reads’ (see **Methods**) that were roughly twice the length of the expected full length reads. **(g)** Tapestation (D5000) of pooled PCR-prepared libraries submitted for ONT sequencing, showing predominant expected product (≈2.2 kb), and minor empty product from p146 (≈900 bp), and lack of product >4 kb corresponding to ‘dimer reads’ seen in (f). **(h)** Table of read counts with valid barcode pairs and valid fraction across libraries originating from plasmids or AAV packaged DNA (different rows), and direct (PCR-free) [central columns] vs. PCR-derived libraries [right columns]. The data from the direct libraries is reproduced from Fig. S3 and S4 for plasmid digest and AAVs respectively.

## METHODS

### Cloning complex libraries of doubly-barcoded AAV constructs with various inserts

To clone a set of doubly barcoded AAV constructs, we digested plasmid CN1839 (Addgene #163509) with PacI & BbsI (NEB), and size selected the 2.9 kb backbone containing the AAV2 ITRs on agarose gel. A GFP stuffer constructs with complex barcodes was created by two steps of PCRs (step 1: primers o924+o925 amplifying GFP from p027, step 2: o926+o927_v2) using Kappa Robust and standard cycling conditions. Random Ns were included in the primers to append random DNA barcodes (15Ns in o926, left BC1; 16Ns in o927_v2 right BC2, **Fig. S1d**) The second PCR was tracked by qPCR and stopped at the inflection point to maintain complexity and limit jackpotting. The barcoded GFP stuffer was then size selected on agarose, and inserted by Gibson assembly (4 uL reaction) in the CN1839 PacI+BbsI digested backbone above. The resulting library was cleaned up (reaction taken to 50 uL with 10 mM Tris 8 buffer, Zymo Clean and Concentrator with 3:1 binding buffer, and eluted in 6 uL water), and 3 uL was electroporated in 25 uL of C3020 cells (NEBs). The resulting complex library (>5 M barcode transformants estimated by plating 0.03%) was grown overnight at 37C and plasmid purified (Qiagen miniprep), generating the parental barcoded plasmid p146 (**Fig. S1b**).

This barcoded ‘dock’ contained two SapI sites internal to the BC1 and BC2 to allow insertion of various libraries of inserts. Just outside the SapI sites, a Nextera read1 adaptor served as homology for Gibson assembly on the left BC1 side (**Fig. S1d**, see below), and another constant homology arm was used on the right BC2 side. Prior to replacing GFP by different classes of inserts, we generated additional dock plasmids to accommodate final constructs of fixed lengths by adding constant inserts outside of the barcoded stuffer. Briefly, p146 was sequentially digested with BglII and PmlI (NEB), the resulting linear product size selected on agarose. Filler sequences of 1291 bp and 1898 bp were generated by PCR amplification from plasmid CN1839 (constant_1291bp with primers o929+o930, constant_1898bp with primers o928+o930) using standard conditions. Following size selection on agarose, these were inserted by Gibson assembly and electroporated as described above, maintaining a high complexity of represented barcodes in the libraries, resulting in barcoded plasmids p147 (with constant_1291bp) and p148 (with constant_1898bp). All parental libraries were confirmed by whole plasmid sequencing (Plasmidsaurus).

Parental barcoded AAV libraries with GFP stuffer flanked by SapI sites:

p146: no additional insert.

p147: with additional constant 1291 bp insert outside of barcodes.

p148: with additional constant 1898 bp insert outside of barcodes.

Six libraries were then constructed to vary the insert length and class (homologous, meaning all components of the library are identical, or non-homologous, meaning that all members of the library are different). To maintain a roughly constant length of 2.3 kb between the AAV ITRs, short inserts were integrated in p148, mid-sized inserts in p147, and long inserts in p146 (**Fig. S1b-c**). All parental plasmids were digested with SapI (NEB) to release the GFP stuffer and the resulting barcoded backbones were size selected on agarose for downstream steps (below).

Homologous (fixed) inserts were taken from sections of the CN1839 cargo (**Fig. S1a**) and generated by two steps of PCR with primers. The first step appended handles (Nextera R1 on left, partial Nextera R2 on right) to the constant region: homologous_127bp with primers o931+o934, homologous_739bp with o931+o933, and homologous_2034bp with o931+o932. These handles then served to prime a secondary PCR which also appended a unique library index inside the construct for later demultiplexing. This secondary PCR was the same as the one used on the construction of the non-homologous libraries, corresponding to primers Nextera R1 (o759) on the forward direction, and an indexed Nextera R2 with the constant right homology arm on the reverse direction (homologous_127bp with o937, homologous_739bp with o938, and homologous_2034bp with o939). The indexed constant inserts were then size selected on agarose, and inserted in their respective (for fixed ITR-to-ITR length) SapI digested barcoded parental backbone (**Fig. S1b**) with Gibson assembly. Following clean up and electroporation, libraries were bottlenecked by serial dilution prior to outgrowth to a target complexity of ∼20k transformants.

To generate non-homologous insert libraries, we relied on tagmentation and PCR amplification of bacterial genomic DNA. Briefly, 1 uL at 10 ng/uL of gDNA extracted from *Escherichia coli* cells (also containing plasmid p146, see below) was tagmented with dually loaded Tn5 (Illumina, Nextera Tagment DNA enzyme, cat. no. 15027916) at two doses: 0.4 uL Tn5 enzyme 1 and 0.4 uL of a 20-fold dilution of the Tn5 enzyme together with 3.6 uL of water and 5 uL of 2x tagmentation buffer (Illumina, cat. no. 15027866). Following clean up (Zymo Clean and Concentrator, 3:1 binding buffer), 1 uL of the 10 µl Tris 8 10 mM elution (1 ng) was taken as input for 12 cycles of PCR (Kappa Robust) from the Nextera handles with indexed primer (same primers series as homologous inserts above, forward o759, reverse: short with o941, mid-sized with o942, and long with o943) to mark the libraries with internal insert indices for downstream demultiplexing. The resulting smear was size-selected to a narrow range in size on polyacrylamide gel for the short insert, and on agarose for the mid-sized and long inserts. In all cases, to size-match non-homologous fragments as carefully as possible, the corresponding length homologous inserts were run on side lanes on the gels and the small corresponding range of the amplified tagmented gDNA was cut out and purified. The size-selected fragments were secondarily amplified with the same primers to generate more material for cloning, size selected again, and inserted by Gibson assembly in their respective SapI digested barcoded parental backbone as for the homologous fragments. Following clean up and electroporation, libraries were again bottlenecked to a target complexity of ∼20k transformants.

All in all, we thus obtained the following six libraries fixed ITR-to-ITR length with the following insert characteristics and estimated complexity from transformant counts. All homologous libraries were confirmed by whole plasmid sequencing (Plasmidsaurus), and non-homologous libraries were spot checked with Sanger sequencing of colonies (Genewiz).

Final dual barcoded AAV libraries with various inserts (listed insert lengths do not include the Tn5 Nextera handles, included between all barcode pairs: 33+34 bp total)

p149: short (127 bp) homologous insert

p150: short (≈100 to 150 bp) non-homologous inserts

p151: mid-sized (739 bp) homologous insert

p152: mid-sized (≈650 to 850 bp) non-homologous inserts

p153: long (2034 bp) homologous insert

p154: long (≈1900 to 2100) non-homologous inserts

We note that the bacterial pellet used for genomic DNA extraction to tagment for non-homologous libraries insert generation was outgrown from a colony on a p146 transformation plate (but grown on LB without ampicillin). As such, inserts from these libraries contained at substantial proportion sequences from plasmid p146 (proportion of mapped fragments: 65% for p150, 19% for p152, 15% for p154, see table on p. 21, inserts mapping to p146 also mapped to CN1839 given similarity). While inserts from these non-homologous libraries are still very diverse, given the limited size of the plasmid and substantial proportions of the libraries, there are still non-zero pockets of homology for certain members of the libraries. To quantify this, we performed local pairwise alignment (pairwiseAlignment from R package Biostrings version Biostrings_2.62.0, option type=“local”) of randomly selected inserts sequences from the size-selected digested plasmid long-reads (pairs of reads with distinct barcodes). Setting a threshold score per library as the maximum of 1000 alignments from a pair of insert sequences and another 1-shuffled pair (FDR<0.001), we quantified that about 1% of inserts had detectable homology (0.8% p150, 1% p152, 1.8% p154), indeed close with theoretical expectation assuming even random fragmentation with the proportion of reads within each library coming from the plasmid (size 5 kb) and the size of inserts [p150: 0.65×0.65×(100 bp/5000bp)=0.8%, p152=0.19×0.19×(750 bp/5000 bp)=0.5%, p154=0.15×0.15×(4000bp/5000bp) = 1.8%]. Hence, despite the tagmentation material in the starting library being a mixture of genome and plasmid, the final libraries were effectively nearly completely non-homologous.

Notably, a significantly enriched proportion of the rare swapped-barcode reads from the non-homologous libraries originated from members of the library with regions of homology. For instance, of the 22 (out of 26) full BC-to-BC length barcode-swapped reads from library p150 for which the two corresponding pre-swap inserts could be mapped in the size-selected digested plasmid data, 3/22 had extensive homology (pairwise local alignment score>50), which was over ten-fold higher than randomly selected pairs of inserts from the same library (Fisher’s exact p<0.005).

### Generation of a valid BC1-BC2 pairs dictionary from parental plasmid library p146

To generate the dictionary of valid barcode 1 and barcode 2 pairs, we amplified by PCR (primers o945+o946v2 containing P5 and P7 Illumina adapters) the GFP stuffer insert flanked by the two barcodes using 5 ng starting template (50 uL reaction, Kapa Robust, standard conditions, 10 cycles) and followed by 1x ampure beads cleanup. The resulting library was sequenced using custom primers as a fraction of a NextSeq2000 P2 100 cycles run (read1: 42 cycles with o947, index1: 20 cycles with o761 Nextera_read1 [into SapI restriction site and GFP, not used], index2: 20 cycles with o948 [into GFP, not used], read2: 16 cycles with o762 Nextera_index2). Sequencing data was demultiplexed from other samples based on the first 10 indices of read1 (GATCCGTCGA) using bcl2fastq with base mask i10y*,y*,y*,y*, yielding 195.6M reads to associate barcodes from p146. BC1 (5’/left of insert) was on cycles 1 to 16 within read 2, BC2 (3’/right of insert) was on cycles 21 to 36 within read1.

We then applied stringent representation and uniqueness criteria to identify unique valid pairs for our downstream swapping assessment. Read counts corresponding to identical barcodes 1 and 2 were first piled up. First, representation of barcodes (separately, summing reads from pairs with 3 or more counts) was inspected (**Fig. S2c**), revealing a trimodal distribution: low counts (<=5 reads) corresponding to likely sequencing or PCR errors (BC1: n=1010349, 2.5% of reads; BC2: n=1162035, 2.9% of reads), intermediate counts barcodes making up the bulk of the coverage (BC1=6345673 barcodes, 92.9% of reads; BC2: n=6596695 barcodes, 93.5% of reads), and high-count barcodes (BC1: n=827, 4.6% of reads; BC2: n=530 barcodes, 3.7% of reads), possibly emerging from of clonal expansion during transformation outgrowth and/or PCR jackpotting. In the case of BC2, a single sequence (ATAACGACTTGTGAGC) was drastically over-represented (2.5% of reads). This barcode and all other barcodes within a Levenshtein distance of 2 or less to it (n=402 barcode, 0.25% of reads) were not considered in downstream analysis to avoid spurious non-unique pairs. We note our barcode space was not saturated at that level of tolerance to mismatches (average of 3 BCs within our BC2 reads within Levenshtein distance of 2 to repeated 1-shufflings of the ATAACGACTTGTGAGC sequence). Further, barcodes with 10 or more consecutive Gs, or containing truncated BC (with the detected post-BC sequences: ATTAAAC for BC1, TAGCGCG for BC2) were removed, corresponding to a minute proportion of the library (BC1: n=9533, 1.1% of reads; BC2: n=3910, 0.06% of reads). To avoid mismatches from high-representation barcodes being retained as spurious mid-counts barcodes, the list of mid-counts barcode was pruned by removing those within Levenshtein distance of 2 from the high-counts barcodes (number mid-counts barcodes removed in pruning process: BC1 n=16386; BC2 n=4747). All remaining pruned mid-counts barcodes were retained downstream (BC1: n=6321002; BC2: n=6588391). Finally, the high-counts barcodes were further error-corrected by generating an undirected graph connecting barcodes within Levenshtein distance of 2 or less. The most highly represented barcodes from each connected component were considered valid. Rare clusters composed of many well-represented barcodes (fold-change between max/min < 10 within connected component) were discarded as possibly ambiguous, leading respectively to n=609 and n=420 error-corrected high count BC1 and BC2 sets. All in all, these filtering steps generated a list of well-represented high quality barcodes (BC1: n=6321611, BC2: n=6588811).

From these well-represented BC1 and BC2, we finally filtered on unique pairing between the two. Specifically, all paired barcodes with 2 or more reads were filtered for members present in both separately valid sets. Then, the proportion of reads to each barcode within a pair (e.g., number of reads to BC1 from a given pair over number of reads containing the same BC1 across all pairs in the library) was computed. Only pairs for which >99% of reads mapped to a unique pair were retained. This led to a final set of n=5586772 valid barcode pairs used for downstream assessment. Only exact matches to BC1 and BC2 constituents of these final pairs were used for filtering long-read data and setting the denominator in our barcode swap quantification. See retention proportion at filtering steps in **Fig. S2b**.

We note that the distinction above between mid- and high-counts representation above was largely immaterial, as the overwhelming majority of detected BC in our long-read libraries were from the mid-counts set (n=12 out of 4811 full BC-to-BC reads from AAV used to quantify swaps in **Fig. 1** from the high-counts barcode set), in line with their dominant representation, with no significant correlation with concordant or discordant pairing and barcode representation class (for the two libraries with detected high-counts barcodes in the final read list, Fisher’s exact: p149: p=0.71; p151: p=0.14).

### AAV packaging

Complex libraries were packaged into PHP.eB or AAV2 capsids using the crude prep method previously described^29^. Maxiprep libraries cloned between AAV2 ITRs were transfected with PEI Max 40K (Polysciences Inc., catalog # 24765-1) into one 15-cm plate of HEK-293T cells (ATCC CRL-11268), along with helper plasmid pHelper (Cell BioLabs), and either pUCmini-iCAP-PHP.eB^30^ (Addgene #103005) or pAAV2/2 (Addgene #104963). The final transfection mix contained 150 μg PEI Max 40K, 30 μg pHelper DNA, 15 μg rep/cap plasmid DNA, and 15 μg library DNA per plate (“Standard conditions”). In the case of “Sparse conditions”, we transfected with fewer library molecules per cell by using 10% (1.5 μg) library DNA along with 90% (13.5 μg) non-ITR-bearing empty expression vector plasmid DNA as a carrier (CN2481, pCDNA3.1-CMV-empty-IRES2-mTFP1-BGHpA). Following transfection at 24 hours the medium was changed to low serum conditions (1% FBS), and then after 5 days cells and supernatant were harvested into 50 mL conical tubes, and AAV particles were released by three freeze-thaw cycles. The cell lysates were then treated with benzonase to degrade free DNA (2 μL benzonase, 30 min at 37°C, MilliporeSigma catalog # E8263-25KU), and then cell debris was cleared with low-speed spin (1500 g 10 min). The supernatant containing virus was concentrated over a 100 kDa molecular weight cutoff Centricon column (MilliporeSigma catalog # Z648043) to a final volume of ∼100 μL containing ∼1-3e12 vector genomes. Crude AAVs were used for direct sequencing (Plasmidsaurus AAV sequencing service).

### Sample preparation & sample submission for long-read sequencing

We long-read sequenced both direct and PCR-based libraries. For PCR-free sequencing of plasmid DNA, we digested the starting dually barcoded AAV plasmid libraries p146 & p149-p154 with NotI-HF and MluI-HF (NEB, using starting material 2 ug, 37C, 1h). The released the barcoded inserts (881 bp for p146, ≈2.3 kb for p149-p154) were then size selected on agarose (Zymoclean gel purification), pooled, and submitted to Plasmidsaurus for a custom long-read project (project ID 8Y6YSQ.1, target 3M reads, recovery 2.6 M reads). PCR-free libraries of AAV-packaged DNA were sequenced using the AAV service from Plasmidsaurus (project ID RP88L6).

To generate libraries of barcoded insert by PCR from the AAV packaged DNA, we first extracted DNA by performing proteinase K treatment (3 uL crude AAV prep, 6 uL 10 mM Tris 8, 1 uL proteinase K [Thermo Scientific EO0491], 60 min at 50C, 5 min at 70C, then placed on ice), followed by phenol/chloroform extraction (add 190 uL 10 mM Tris 8, add 200 uL phenol-chloroform-isoamyl alcohol [Invitrogen 15593-031], vortex 30 s, spin at 16,000 g at room temperature for 5 min, take aqueous layer) and isopropanol precipitation (add 1 uL glycoblue, 50 uL NaOAc 3M, 250 uL isopropanol 100%, vortex, 45 min at -80C, 45 min at 21,000 g at 4C, 80% EtOH wash, air dry pellet, resuspend in 10 uL 10 mM Tris 8). 2 uL of the precipitated DNA was taken as template for PCR with primers oJBL949+oJBL950 (Kapa Robust Hot Start Ready mix [Roche], 60C annealing temperature, elongation time 2 min 30 s) and tracked by qPCR (SYBr green). PCR libraries were generated from all samples packaged with the PHP.eB serotype in standard (non-sparse) packaging conditions (i.e. samples 1 to 6, see Fig. S4f). Two separate reactions per sample (reaction 1: volume 20 uL, 16 cycles; reaction 2: volume 50 uL, 17 to 20 cycles) were pooled prior to submission. Of note, we tested the importance of adding a DNase treatment step by performing qPCR with primers targeting the insert between the AAV ITRs (oJBL905+oJBL906) vs. a sequence on the ampicillin cassette in the backbone (oJBL097+oJBL098) comparing with/without DNase treatment prior to proteinase K treatment but saw no difference (>30-fold lower backbone material relative to cargo), likely because of the benzonase treatment already having degraded the non-encapsidated DNA. PCR libraries from plasmids were prepared from 1 ng of template (with quantification possibly confounded by genomic DNA carryover), and reactions were stopped at 18 cycles (starting input material calibrated to still be in exponential phase before the inflection point). Plasmid-templated samples also included the parental p146 (which was pooled at 1/10-fold stoichiometry of other larger inserts to mitigate ONT size biases). In both plasmid- and AAV-templated conditions, PCR reactions were cleaned up with 0.4x Ampure clean, leaving predominantly >2 kb products, **Fig. S5g** (i.e., largely excluding p153^×^-short and p153^×^-mid samples). Two samples corresponding to pooled products (AAV pool sample 1, plasmid pool sample 2) were submitted to Plasmidsaurus for long-read sequencing (custom project ID: DXYB6C, target reads 1M per sample).

### Long-read sequencing data analysis

ONT reads were processed by first searching for ‘signpost’ sequences (**Fig. S1d, S3b**), corresponding to constant regions flanking barcodes, within the reads.

Signpost sequences:

Left BC1 upstream: CGCGTTTAAT

Left BC1 downstream: TCGTCGGCAGCGTCAGATGTGTATAAGAGACAG

Right BC2 upstream: GATCCGTCGATCGACTGAGT

Right BC2 downstream: TAGCGCGCCATTAGGCATGA

To do so, we ran Smith-Waterman (ssw-align^31^, option -r) using the ONT reads as target and signposts as query, and reformatted the output to collate alignment scores (**Fig. S3c, S4b**) from all signposts for each target read. Reads for which all 4/4 signposts were detected (75% match or higher) in the concordant orientations and expected barcode order (BC1 position < BC2 for + strand, and vice versa) were retained. The sequences corresponding to barcodes, insert index (one for each library), and insert were obtained based on local position reference frames set by signpost positions and length and strand. All sequences were then converted for forward strand for downstream analysis (reverse complement for reads mapped to - orientation). Reads were demultiplexed based on the insert index by calculating the Levenshtein distance to the closest index from the expected list:

| Insert class | Insert index | Indexed primer used for insert amplification |
| --- | --- | --- |
| p146 (parental) | CTCTTCAGAT | NA |
| Short, homologous | AGACTCAAGT | o937 |
| Short, non-homologous | GAGACGGTCA | o941 |
| Mid-sized, homologous | AGACGATCTG | o938 |
| Mid-sized, non-homologous | GAGACCCTAA | o942 |
| Long, homologous | AGACATTGGC | o939 |
| Long, non-homologous | GAGACCATCG | o943 |

The second best match was also retained. Any reads for which the best match was within a Levenshtein distance 1, and the difference to the second best match was at least 2 was deemed unambiguously demultiplexable (**Fig. S3d, S4d**). Finally, only reads with BC1 and BC2 (separately, i.e., irrespective of whether their pairing is correct) corresponding to exact match in our final set of barcodes in our dictionary of valid pairs were retained from quantification. Reads were deemed ‘full length’ with regards to BC-to-BC distance if they fell within the 1% to 99% size distribution from the size selected & digested reads (demarcated by vertical lines in the BC-to-BC length distributions of **Fig. S3e**, see table in **Fig. S3g**). Only full BC-to-BC length reads were considered for quantification in **Fig. 1c-d**, see table in **Fig. S4f** for proportion of valid barcode pairs from reads outside of the range in long-insert libraries (‘non full length’), which contained a high proportion of shorter reads (**Fig. S4d**). See **Fig. S3a** and **S4a** for the tabulation of the number of retained reads at each filtering step from plasmid-digested inserts and AAV-packaged material respectivley. Full ITR-to-ITR length boundaries for classification as full length read (analysis of **Table S3**) were taken to be: plasmids parental p146: 800 to 1050 nt; plasmids all others: 2000 to 2500 nt, AAV all except p153^×^-mid and p153^×^-short: 2250 to 2750 nt, AAV p153^×^-mid: 1100 to 1400 nt, and AAV p153^×^-short: 500 to 750 nt. These boundaries are demarcated by vertical lines in the read length distributions of **Fig. S3f** and **S4e**.

To collate metadata on insert sequences, we aligned the inserts to both the *E. coli* genome (GCF000005845.2) and plasmid CN1839 (Addgene #163509) using minimap2^32^ (version 2.28, option -x map-ont). Alignments with >0.65 fraction of matching bases (matching bases in alignment/alignment length including gaps) were retained. For a given read and start alignment position within the read, the best scoring (fraction of matching bases) alignment was retained. To further reduce redundant alignments to homologous sections, each read with multiple listed alignments was considered. Overlap (in the read reference frame) between alignments was computed, and alignments were considered redundant if they overlapped by >50% (in practice, the distribution of overlap was nearly perfectly bimodal between 0% and 100%), and the best alignment was retained. To properly categorize inserts overlapping with the end of the plasmid (minimap2 does not deal with circular chromosomes), reads with two disjoint alignments and one starting within position 5 nt or ending within 5 nt of the end of the plasmid were re-categorized as non-composite (unique overlapping the circular junction). Based on these, read inserts were then categorized on whether they contained a non-mapped (based on our stringent procedure), a unique alignment (to either the *E. coli* genome or CN1839), or a composite alignment (two or more non-overlapping alignments to either the *E. coli* genome, CN1839, or both). Quantification (table below) from the plasmid inserts aligned with expectations (homologous libraries entirely comprised of simple fragments mapping to the plasmid, non-homologous libraries coming from a mixture of both given the source material used for tagmentation, see p. 17).

Fraction of alignment categories for inserts of size-selected digested plasmid reads:

| Plasmid ID | simple<br>CN1839 | simple<br>*E. coli* | not mapped | composite<br>CN1839 | composite<br>*E. coli* | composite<br>*E. coli*/CN1839 |
| --- | --- | --- | --- | --- | --- | --- |
| p146 (parental) | 0.9954 | 0.0000 | 0.0042 | 0.0002 | 0.0000 | 0.0002 |
| p149 (short, homologous) | 0.8434 | 0.0002 | 0.1551 | 0.0008 | 0.0000 | 0.0005 |
| p150 (short, non-homologous) | 0.6083 | 0.3212 | 0.0694 | 0.0005 | 0.0002 | 0.0005 |
| p151 (mid-sized, homologous) | 0.9003 | 0.0006 | 0.0975 | 0.0010 | 0.0001 | 0.0005 |
| p152 (mid-sized, non-homologous) | 0.1889 | 0.7853 | 0.0179 | 0.0002 | 0.0058 | 0.0019 |
| p153* (long, homologous) | 0.9962 | 0.0000 | 0.0028 | 0.0008 | 0.0000 | 0.0003 |
| p154 (long, non-homologous) | 0.1472 | 0.8216 | 0.0158 | 0.0018 | 0.0126 | 0.0010 |

Detailed insert alignment metadata is included in the final tables of reads, and compiled for statistical comparison in full length BC-to-BC vs. non-full length BC-to-BC reads, in which simple vs. composite/non-mapped reads were compared for each library (Fisher’s exact test) in **Table S2**, with incomplete BC-to-BC reads being composed of a higher proportion of composite reads.

For the PCR derived libraries, a small subset (variable proportion across libraries) of reads had roughly twice the length of the expected full ITR-to-ITR products (**Fig. S5f**, grey arrows). Inspection of these longer reads revealed that they corresponded to both a forward and a reverse pass of the same sequence. Given that submitted products were predominantly of the expected size (≥95%, **Fig. S5g**), we tentatively attribute these to self-ligation during the end-repair & adapter ligation step in the ONT library prep. Indeed, the junction of the forward/reverse portion of these reads corresponded to the near perfect hairpin sequence, GCGGACCGAGCGGCCGC, which might have created a substrate for hairpin ligation to the other strand. Regardless, the frequency of barcode swaps from normal reads vs. these ‘dimer’ reads were not substantially different across all libraries (Fisher exact test p>0.2 in all cases except AAV p153 long homologous, for which p=0.007 but with fraction discordant dimer reads equal to 0.56 in the dimer reads vs. to 0.49 in the non-dimer reads and were thus also quite similar).

### Statistical testing (bootstrap false discovery rate)

To provide estimates of significance from counting noise (some AAV samples had relatively few reads passing quality control filters, **Fig. S4f**), bootstrap resampling was performed to generate ensemble estimates of concordant barcode pairs. To compare two samples (e.g., p149:AAV vs. p150:AAV in Fig. 1c), n=10^5^ bootstrap re-samplings were performed. In this case, the bootstrap false discovery rate (FDR) was taken as the fraction of re-samplings in which sample p149:AAV had higher bootstrap concordant pair fraction than the p150:AAV re-sampling, etc.

